# Temporal Phase Differences Encode Tactile Motion from Static Vibrotactile Inputs

**DOI:** 10.64898/2026.01.24.701470

**Authors:** Erfan Rezaei, Mehdi Adibi

## Abstract

Motion perception in touch is traditionally attributed to spatially sequential stimulation or skin deformation caused by moving objects. Here, we demonstrate that directional tactile motion can instead be inferred purely from temporal phase differences between two spatially static vibrotactile inputs. Using psychophysical experiments in human participants (n = 56), we show that continuous vibrations delivered to separate fingertips, when offset in phase, evoke a robust and directionally consistent motion percept despite the absence of physical movement. Discrimination performance depended systematically on the temporal phase relationship between stimuli, following a sinusoidal relationship, consistent with a phase-based motion inference mechanism. Vibration amplitude had little influence on perceptual accuracy once stimuli exceeded detection threshold, although higher amplitudes modestly reduced response times. In contrast, envelope dynamics and spatial configuration significantly modulated performance: vibrations with exponential amplitude envelopes – reflecting the natural propagation of mechanical energy through a medium – enhanced phase sensitivity and sped responses, and bimanual stimulation across the two hands improved both accuracy and response time. These findings identify temporal phase integration as a fundamental mechanism for tactile motion perception, analogous to timing-based computations in auditory localisation and visual motion detection. Phase-coded vibrotactile stimulation therefore offers a minimal and efficient strategy for conveying directional motion in haptic interfaces without requiring spatial arrays or moving actuators.

## Introduction

Perception of tactile motion is integral to object manipulation, surface exploration, texture perception, and spatial awareness (Johansson and Flanagan, 2009; Lezkan and Drewing, 2018; Boundy-Singer et al., 2017; Weber et al., 2013; Augurelle et al., 2003; Adibi, 2025). During active touch, tactile motion cues provide critical information about object geometry and surface structure, supporting both exploratory and manipulative behaviours (Johansson and Flanagan, 2009; Janko et al., 2018). Beyond natural tactile motion arising from physical moving contact (Olausson and Norrsell, 1993), tactile motion percepts can also be evoked through spatiotemporal modulation of pulsed or continuous vibrotactile stimuli delivered to static skin locations (Sherrick and Rogers, 1966; Kirman, 1974; Gardner and Palmer, 1989; Pei and Bensmaia, 2014; Adibi, 2025). Classic psychophysics studies established that sequential stimulation at two loci (e.g., along the arms or on the back) generates reliable apparent tactile motion, where perceived direction and continuity depend strongly on inter-stimulus timing and stimulus duration (Sherrick and Rogers, 1966; Kirman, 1974). Related spatiotemporal tactile illusions, such as the cutaneous rabbit illusion, demonstrate that sequences of brief stimuli delivered at separated skin sites can produce percepts that interpolate between stimulation sites (Geldard and Sherrick, 1972). This highlights that temporal structure is a key element in constructing spatially organised percepts.

In our previous study (Adibi, 2025), we extended this framework to continuous and smooth perception of motion across fingertips. This paradigm differs from conventional apparent motion paradigms applied to skin regions with relatively lower tactile acuity, such as the arms or back, or within a single finger using pulsed stimulation and dense arrays of tactors. Specifically, we demonstrated that two continuous amplitude-modulated vibrotactile stimuli delivered to separate fingertips, with a controlled phase offset in their modulation envelopes, generate robust continuous motion percepts. We further analysed two classes of computational models that map spatiotemporal vibrotactile input to perceived motion by exploiting different types of stimulus evidence. One class is based on the spatiotemporal correlation structure between signals presented at different skin locations. In these models, motion direction emerges from asymmetric correlations produced by temporal delays between spatial channels, consistent with a delay-and-correlate computation (e.g., the Hassenstein–Reichardt detector) and closely related motion-energy models widely studied in visual motion perception (Hassenstein and Reichardt, 1956; Adelson and Bergen, 1985; Borst, 2000). In contrast, landmark-based models posit that motion direction is inferred from the relative timing of salient temporal features in the stimulus envelope (such as peaks, troughs, or points of maximal slope). In this framework, motion estimation precision depends on how sharply these landmarks are defined, leading to predictions that vary with the envelope waveform and feature salience (Adibi, 2025). Critically, these two computational frameworks can make overlapping predictions for some stimulus families (e.g. smooth symmetric envelopes), but diverge when stimulation features such as envelope asymmetry, sharpness, or timing manipulations differentially affect correlation structure and feature timing (Adibi, 2025).

In the present study, we conducted a series of psychophysics experiments to characterise how systematic manipulations of vibrotactile stimulus parameters shape perceived tactile motion. We further tested how these effects differentiate the predictions of correlation-based and landmark-based computations. Experiment 1 examines the effect of modulation amplitude on motion perception. Both model classes predict that, once the stimulus is reliably detectable, modulation amplitude has minimal influence on directional accuracy. Experiment 2 provides a critical test of waveform-dependent predictions. Correlation-based models predict invariance to modulation envelope waveform, whereas landmark-based models predict improved discrimination when the envelope contains sharper temporal features. Exponential envelopes are of particular interest because they approximate the natural decay of vibratory energy with distance during physical motion across a surface (Adibi, 2025) and provide more temporally localised landmarks than sinusoidal modulation.

Experiments 3 and 4 extend this framework by examining how temporal motion cues are integrated across spatial configurations, including within-hand fingertip pairings and bimanual stimulation. Although neither model makes specific quantitative predictions for these manipulations, previous work on tactile spatial integration and bilateral processing suggests that anatomical configuration and hemispheric distribution can influence the efficiency with which temporally structured tactile information is combined (Craig, 1985; Tamè et al., 2017; Kuroki et al., 2017; Arslanova et al., 2022). Together, these experiments provide a systematic assessment of how spatiotemporal stimulus structure and spatial configuration shape tactile motion perception, and how well alternative model classes account for these effects.

## Materials and Methods

Four psychophysics experiments were conducted to investigate vibrotactile motion direction discrimination using a common two-alternative forced-choice (2AFC) discrete-trial paradigm, following the general procedure described in (Adibi, 2025). Briefly, on each trial, participants reported the perceived direction of motion (left versus right) generated by two simultaneously delivered amplitude-modulated (AM) vibrations applied to two fingertips. Responses were made by pressing arrow keys (Experiment 2 and 3), or one of two foot switches (Experiment 1 and 4) with the corresponding foot (left or right) indicating the perceived direction of motion. Trials were self-paced, and participants were able to respond at any time during or after stimulus presentation. Experimental conditions were pseudo-randomised within each session such that participants could not predict the upcoming condition. For Experiments 1, 2, and 4, each stimulus condition was repeated 60 times per participant, with equal probability of leftward and rightward motion. All experimental procedures were approved by the Monash University Human Research Ethics Committee (MUHREC).

### Participants

Fifty six healthy adults (39 female; age range 19–43 years) participated across the four experiments. The same participants took part in Experiment 1 and 4. All participants reported normal tactile perception and no history of neurological disorders. Written informed consent was obtained from all participants prior to participation.

### Apparatus and stimuli

The apparatus and general stimulation procedures were identical to those described in our previous study (Adibi, 2025). Briefly, vibrotactile stimulation was delivered using miniature solenoid actuators (Tymphany PMT-20N12AL04-04; 4 Ω, 1 W, 20 mm nominal diameter), mounted on a vibration-isolated platform. Depending on the experiment, two, three, or four solenoids were used (see below).

Vibrotactile stimuli were generated in MATLAB (MathWorks, Natick, MA; sampling rate of 4.1 kHz) and delivered via multichannel sound cards (Creative Sound Blaster Audigy Fx or Creative Sound Blaster Live! 24-bit External, SB0490). In Experiments 3 and 4, actuators were driven using a 3 W stereo amplifier module (PAM8403 chipset). In all experiments, actuator outputs were matched across channels and configurations. Sound pressure level (SPL) was measured 5 mm from each solenoid and adjusted to match levels across actuators (mean SPL of 78.93 dB). Levels were further verified perceptually to ensure comparable subjective strength across channels.

Vibrotactile stimuli consisted of a 100 Hz carrier sinusoid amplitude-modulated at 0.5 Hz, with a maximum duration of three modulation cycles (6 s). Motion percepts were generated by a fixed phase offset (*φ*) between the modulation envelopes of simultaneously active actuators. All trials began at an iso-amplitude point of the modulation envelope. In Experiment 2, modulation envelope waveform was varied (sinusoidal versus exponential; see below).

For Experiments 1, 2, and 4, solenoids were mounted with a fixed centre-to-centre spacing of 5 cm. In Experiment 3, solenoid positions were adjusted individually for each participant to accommodate differences in finger length and spacing and to ensure comfortable contact with the index, middle, and ring fingertips of the right hand.

### Experimental Conditions

Four experiments characterised vibrotactile motion perception for different stimulus configurations and parameter manipulations.

In Experiment 1, stimulation was delivered to the index and middle fingertips of the right hand using sinusoidal modulation envelopes. Phase offsets of *±*30°, *±*60°, and *±*90° were tested at two modulation amplitude levels – referred to as ‘low’ and ‘high’ – to quantify the effect of envelope amplitude on motion direction discrimination. The low-amplitude condition was set to half the amplitude used in the high-amplitude condition, with the high-amplitude level corresponding to that used in the other experiments (n=16 participants).

Experiment 2 used the same unimanual two-fingertip configuration (right index and middle fingertips), but compared sinusoidal modulation envelopes with exponential envelopes defined as

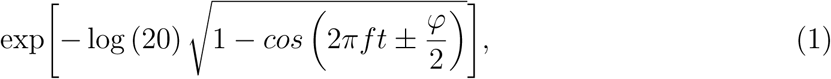

where *f* denotes the envelope frequency (0.5 Hz) and *φ* denotes the phase offset between envelopes. Phase offsets ranged from *±*30° to *±*150° in 30° increments (*n* = 8 participants).

In Experiment 3, three actuators were positioned under the index, middle, and ring fingertips of the right hand, allowing comparison of adjacent (index–middle, middle–ring) and non-adjacent (index–ring) fingertip pairings. Phase offsets were adaptively varied be-tween 5° and 60° in 5° increments using a QUEST adaptive staircase procedure (Watson, 1983), targeting 70% correct performance. This resolution is above the temporal resolution limit imposed by the 100 Hz carrier (10 ms per cycle), which corresponds to approximately 1.8° of the 0.5 Hz modulation cycle, while providing sufficiently fine-grained sampling of phase differences and maintaining stable and efficient convergence of the adaptive procedure. Each fingertip configuration comprised 50 trials, randomly interleaved within a session, yielding a total of 150 trials per participant.

The psychometric function was modelled as a Weibull function, with an initial threshold estimate of 7.5° and a prior standard deviation of 15°, slope parameter of 0.946, lapse rate of 0.06, and guess rate of 0.5. These parameter values were derived from group-level pilot data in Adibi (2025). Threshold estimates were defined as the posterior quantile of the threshold parameter (Pelli, 1987).

Seventeen participants were included in the analysis, defined as those for whom the QUEST procedure converged in at least one of the three fingertip configurations. For one participant, the staircase did not converge in the middle-ring (MR) condition. Data from that condition were excluded, resulting in *n* = 17 for index–middle (IM) and index–ring (IR), and *n* = 16 for MR.

Phase offsets proposed by QUEST were rounded to the nearest multiple of 5° within the range 5°–60°, and the direction of motion (left or right) was randomly assigned with equal probability on each trial. When QUEST proposed phase offsets that exceeded 60°, values were truncated at this upper limit to prevent convergence to values near 180°, where motion perception becomes ambiguous and performance returns to chance (50%), resulting in dual threshold solutions (Adibi, 2025). The prior behavioural results and model predictions indicate a peak discrimination performance around 60°–90° (Adibi, 2025).

Because thresholds beyond 60° cannot be reliably quantified, threshold estimates were treated as right-censored at 60°. Censoring occurred in 3 of 16 participants for the MR configuration, and in 3 and 2 of 17 participants for IM and IR configurations, respectively. Accordingly, performance was characterised using two complementary measures: (i) empirical cumulative distribution functions (ECDFs) of QUEST threshold estimates within the 5°–60° range, and (ii) discrimination accuracy at the maximum tested phase offset (60°), summarised using rank-based statistics across participants for each fingertip configuration.

Differences between ECDFs were assessed using within-subject permutation tests on the area between cumulative distributions (10,000 permutations). For each pairwise comparison, the configuration labels were randomly permuted within participants, preserving the within-subject structure, and the area between the ECDFs was used as the test statistic. Statistical significance was assessed using two-tailed tests. Post hoc power analyses were conducted using G*Power version 3.1.9.7 (Faul et al., 2007) to estimate the sample sizes required to detect the observed effect sizes in pairwise comparisons (two-tailed Wilcoxon signed-rank tests, *α* = 0.05, power = 80%).

Experiment 4 characterised the effect of spatial configuration by comparing unimanual and bimanual stimulation. Four actuators were mounted under the index and middle fingertips of both hands, although only two actuators were active on any given trial. Motion was delivered either unimanually (index and middle fingertips of the right hand) or bimanually (index fingertips of both hands). Phase offsets of *±*30°, *±*60°, and *±*90° were tested (*n* = 16 participants).

### Landmark Model of Motion Perception

A plausible computation underlying tactile motion perception is the landmark model (Adibi, 2025) in which the tactile system detects specific temporal landmarks in the envelope of each vibration (such as peaks, troughs, or other salient features) and infers the motion direction based on the temporal order or timing of these events relative to each other. Here, we assume that a point of reference (or landmark) is detected when the change in the vibration envelope exceeds a certain slope or amplitude threshold. Changes below this threshold are not perceived as distinct events. Thus, any portion of the envelope around the true landmark whose amplitude lies within the threshold margin is perceptually in-distinguishable from the true peak, resulting in temporal uncertainty in the perceived moment of the landmark. An inaccurate motion direction, therefore, is due to overlap between the temporal uncertainty windows of one vibration relative to the other vibration, as formulated in detail in (Adibi, 2025).

### Statistical Analyses

Response accuracy and reaction time (RT) were quantified at the participant level for each experimental condition and phase offset. Accuracy was defined as the proportion of correct trials. RT was defined as the time elapsed between stimulus onset and response time.

Trials were excluded on a within-participant basis if RTs were less than 150 ms (anticipatory responses) or exceeded the participant’s mean RT by more than 2.5 standard deviations. The mean and standard deviation were calculated across all trials for each participant. This filtering removed less than 2.3% of the trials (*n* = 309 out of 13,440) across all participants. The resulting filtered dataset was used for all subsequent analyses, including descriptive summaries and mixed-effects modelling.

Condition-level summaries were computed by averaging these measures across trials within each participant, and group-level summaries were obtained by averaging across participants. Values were reported as mean *±* SEM (standard error of the mean), unless otherwise specified.

For each condition, 95% confidence intervals (CIs) for group means were estimated using a participant-level nonparametric bootstrap (resampling participants with replacement). Percentile-based CIs were obtained from the 2.5th and 97.5th percentiles of the bootstrap distribution of the mean. Condition effects (low-versus high-amplitude, sinusoidal versus exponential envelope, and bimanual versus unimanual configuration) were quantified using paired within-participant difference scores, with CIs obtained by boot-strapping these differences across participants. For sample sizes *>* 10, CIs were estimated using 10,000 bootstrap resamples, while for sample sizes *≤* 10, CIs were derived from the exact bootstrap distribution of the mean by exhaustive enumeration of all possible re-sampling configurations. Reliability was assessed using 95% bootstrap CIs of the paired differences, with effects considered reliable when the CI excluded zero. In addition, statistical significance at the trial level was evaluated using generalized linear mixed models (GLMMs) at *α* = 0.05, from which regression coefficients, standard errors, confidence intervals, and p-values are reported.

Accuracy and RT were further analysed at the individual trial level using generalised linear mixed models (GLMMs) accounting for the hierarchical structure of the data (trials nested within participants). Accuracy was modelled using a binomial distribution with a probit link. Fixed effects included the sinusoidal phase term sin(*φ*), capturing the periodic and directional nature of phase differences, and its interaction with the relevant experimental factor (amplitude, envelope waveform, or hand configuration). Participant-specific random slopes were included for sin(*φ*) to account for variability in phase sensitivity across participants. Model structures with and without main effects of the experimental factors were compared. Models excluding these main effects and including only interactions with sinusoidal phase term sin(*φ*), provided better fits and avoided implausible baseline shifts at 0° and 180° phase offsets (Adibi, 2025).

Consequently, the experimental factors were included only through interactions with the sinusoidal phase term, allowing their effects to be interpreted as modulations of phase sensitivity rather than baseline performance.

RTs were analysed using generalised linear mixed-effects models (GLMMs) with either Gamma or Inverse Gaussian distributions and a log link, reflecting the positive skewness and non-negativity of RT data. Model comparison based on the Akaike Information Criterion (AIC) and the Bayesian Information Criterion (BIC) indicated that the Gamma distribution provided a substantially better fit than the Inverse Gaussian in Experiments 1 and 4 (AIC/BIC differences of approximately 500 units). In Experiment 2, the Inverse Gaussian distribution provided the best fit (ΔAIC = *−*381, ΔBIC = *−*381).

Fixed-effects structures were further compared within each experiment. In Experiment 2 and 4, the reduced models excluding the main effects of experimental factors and including only their interactions with the sinusoidal phase term sin(*φ*) provided the best fit (Experiment 2: ΔAIC = *−*397, ΔBIC = *−*397; Experiment 4: ΔAIC = *−*2, ΔBIC = *−*9).

In Experiment 1, the best-fitting RT model included the main effects of sin(*φ*) and the envelope waveform, resulting in lower AIC and BIC values compared to a model that additionally included their interaction (ΔAIC = *−*2, ΔBIC = *−*9).

## Results

In our previous study (Adibi, 2025), we demonstrated that continuous spatially static, amplitude-modulated vibrotactile stimulation delivered to two fingertips can evoke a robust percept of directional motion when stimuli differ in temporal phase in their modulation envelope. This established phase-difference coding as a sufficient and robust mechanism for tactile motion perception in the absence of physical skin movement. The present study extends those findings by systematically characterising how key stimulus and spatial parameters shape this phase-driven motion percept. Across a series of psychophysical experiments, we examined the effects of modulation amplitude (Experiment 1), envelope waveform (Experiment 2), fingertip pairing within a hand (Experiment 3), and spatial configuration across hands (Experiment 4) on both discrimination accuracy and response time. Together, these experiments delineate the parametric constraints and integrative properties of temporal phase-based tactile motion perception.

### Experiment 1: Effect of Modulation Amplitude

Both the correlation-based and landmark-based models of tactile motion perception established in our previous study (Adibi, 2025) predict that directional discrimination should depend primarily on the relative temporal phase between stimuli, rather than on the absolute modulation amplitude, once afferent activation exceeds detection threshold. In correlation-based frameworks, motion inference is invariant to envelope magnitude by construction, whereas landmark-based models predict only modest amplitude-dependent effects arising from small shifts in the timing due to changes in the proportion of subthreshold signal segments relative to landmarks. In Experiment 1, we quantified how modulation amplitude influences phase-based tactile motion discrimination by assessing its effects on both perceptual accuracy and response time.

To characterise the influence of stimulus intensity on tactile motion perception, we compared discrimination accuracy and response time for sinusoidally amplitude-modulated vibrations presented at two amplitude levels differing by a factor of two – referred to as ’high’ and ’low’. In both conditions, motion direction was determined by the temporal phase differences between two spatially static vibrotactile stimuli delivered to the index and middle fingers of the right hand.

Discrimination accuracy depended on the temporal phase relationship between the two vibrations, consistent with a sinusoidal dependence on phase difference (Adibi, 2025), but was largely invariant to the modulation amplitude (Figure 1A and B). Across participants and phase differences, the average difference in accuracy between low- and high-amplitude conditions was 1.3% *±* 1.1% (mean *±* SEM), despite the twofold increase in stimulus amplitude. Small amplitude-related differences were observed at individual phase offsets, with the largest difference occurring at 30° (only 4.3% *±* 1.8%), whereas differences at 60° and 90° were negligible (*<* 1%) and not statistically significant (95% bootstrap CIs included zero).

**Figure 1:**
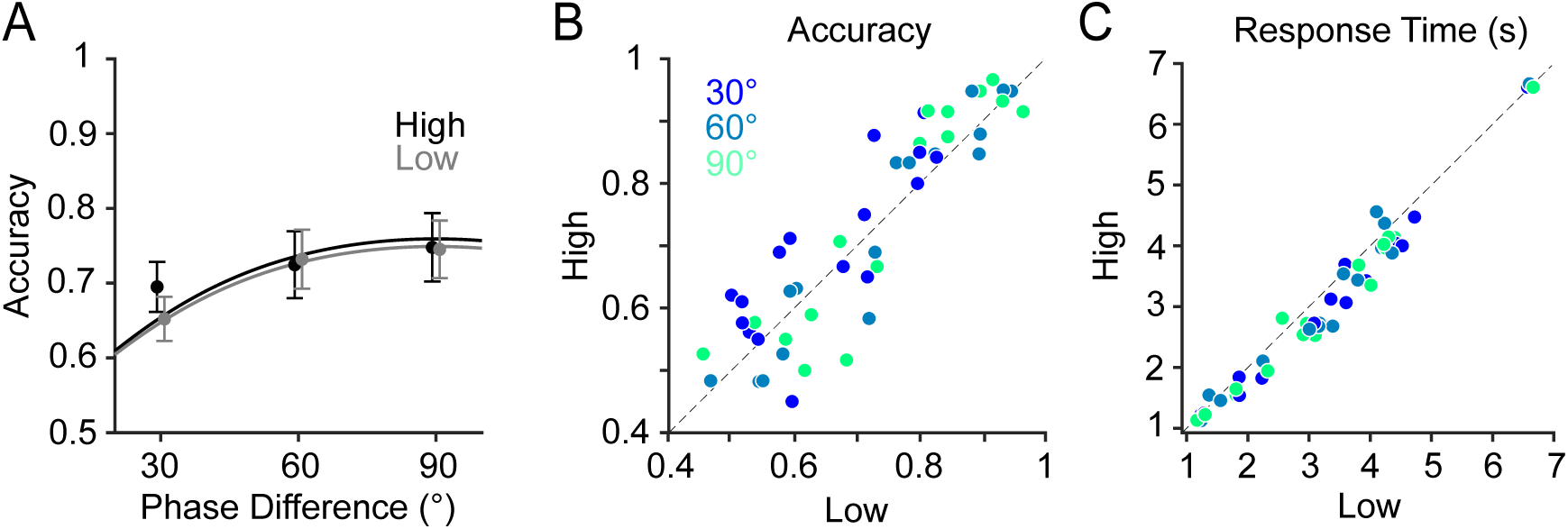
Effect of modulation amplitude on tactile motion perception (Experiment 1). (A) Mean motion direction discrimination accuracy (proportion correct) as a function of phase difference for low- and high-amplitude conditions. Markers indicate mean values and error bars represent standard error of the mean (SEM) across participants (*n* = 16). Curves represent population-level predictions from the GLMM, obtained by averaging subject-level conditional predictions across participants. (B) Accuracy in the high-amplitude condition versus accuracy in the low-amplitude condition. Each data point represents a participant–phase combination and is colour-coded by phase offset (30°, 60°, and 90°). The diagonal dashed line indicates equality between conditions. (C) Same as (B), but for response times.

A hierarchical analysis of accuracy at the single-trial level, accounting for cross-subject variability, confirmed these observations. Accuracy data were analysed using a binomial generalised linear mixed-effects model (GLMM) with a probit link function. The model included fixed effects of the sinusoidal phase-offset evidence term sin (*φ*) and its interaction with the modulation amplitude, with random slopes capturing subject-specific variability in sensitivity. The term sin (*φ*) was a significant predictor of response accuracy (*β* = 0.79 *±* 0.15, 95% CI: [0.5, 1.08], *p <* 10^−6^) indicating robust accumulation of phase-dependent tactile evidence. In contrast, the effect of the modulation amplitude was approximately 20 times smaller and did not reach significance (*β* = +0.04 *±* 0.05, 95% CI: [-0.05, 0.13], *p* = 0.42). Model diagnostics indicated an adequate fit to the data (Figure 1A) with no evidence of overdispersion beyond the binomial assumption (dispersion parameter of 1.001, *n* = 5,594).

Response time provides a behavioural index that is often interpreted as reflecting the strength of the perceived motion signal (Adibi, 2025). We characterised the extent to which modulation amplitude influences response time during motion direction discrimination. Across participants and phase differences, high-amplitude stimulation was associated with consistently faster response times, with an overall mean reduction of 0.22 s. On average, across participants, the response times were significantly faster by 0.26 *±* 0.05 s at 30°, 0.17 *±* 0.07 s at 60°, and 0.22 *±* 0.05 s at 90° (Figure 1C), with 95% nonparametric bootstrap CIs of the response time differences excluding zero at all phase offsets.

A hierarchical analysis of response times at the single-trial level accounting for cross-subject variability, confirmed these observations. Response times were modelled using a Gamma GLMM with a log link function. Fixed effects included modulation amplitude and sinusoidal phase-difference evidence term sin(*φ*). Response times decreased significantly with phase difference (*β* = *−*0.079 *±* 0.031, 95% CI: [-0.141, -0.018], *p* = 0.011) and responses were modestly faster in the high-compared to the low-amplitude condition (*β* = *−*0.077 *±* 0.013, 95% CI: [-0.103, -0.051], *p <* 10^−9^). Model diagnostics indicated an adequate fit, with no evidence of excessive residual variability under the Gamma model (estimated square-root dispersion = 0.497, dispersion = 0.247; *n* = 5,594 trials).

### Experiment 2: Effect of Modulation Envelope Waveform

Experiment 2 quantifies how the temporal profile of the amplitude modulation envelope influences tactile motion perception. As outlined earlier, correlation-based models of motion perception predict invariance to modulation waveform once the signal exceeds vibration detection threshold, whereas landmark-based models predict enhanced directional sensitivity for envelopes with sharper temporal features. To test these competing predictions, we compared motion discrimination accuracy for sinusoidal modulation versus exponential envelopes which approximate the natural profile of vibratory intensity during physical motion (Adibi, 2025).

Accuracy depended on both temporal phase difference and modulation envelope waveform (Figure 2A and B). Across tested phase offsets, exponential vibrations consistently resulted in higher motion direction discrimination accuracy than sinusoidal envelopes: At 60°, accuracy for exponential envelope was 84.2% *±* 5.2%, exceeding performance for sine envelope (77.3% *±* 6.5%) by 6.9% *±* 1.3%. A similar trend was observed at 90°, where accuracy reached 89.3% *±* 4% for the exponential envelope compared to 82.0% *±* 5.5% for the sine envelope (+7.3% *±* 1.8%). Performance differences remained evident at larger phase offsets, with exponential envelopes outperforming sine envelopes by 6.1% *±* 2% at 120° and 4.9% *±* 2.5% at 150°. These differences were statistically significant at all phase offsets except 30°, with 95% nonparametric subject-level bootstrap CIs of differences excluding zero. When averaged across phase offsets (30°–150°) within each participant, exponential envelopes showed a net improved accuracy of 5.3% *±* 1.5% (mean *±* SEM across participants). Notably, despite this systematic advantage, participants still demonstrated robust motion discrimination with sinusoidal envelopes, confirming our previous findings that directional cues can be reliably extracted from phase differences, even in the absence of physically naturalistic envelope dynamics (Adibi, 2025).

**Figure 2:**
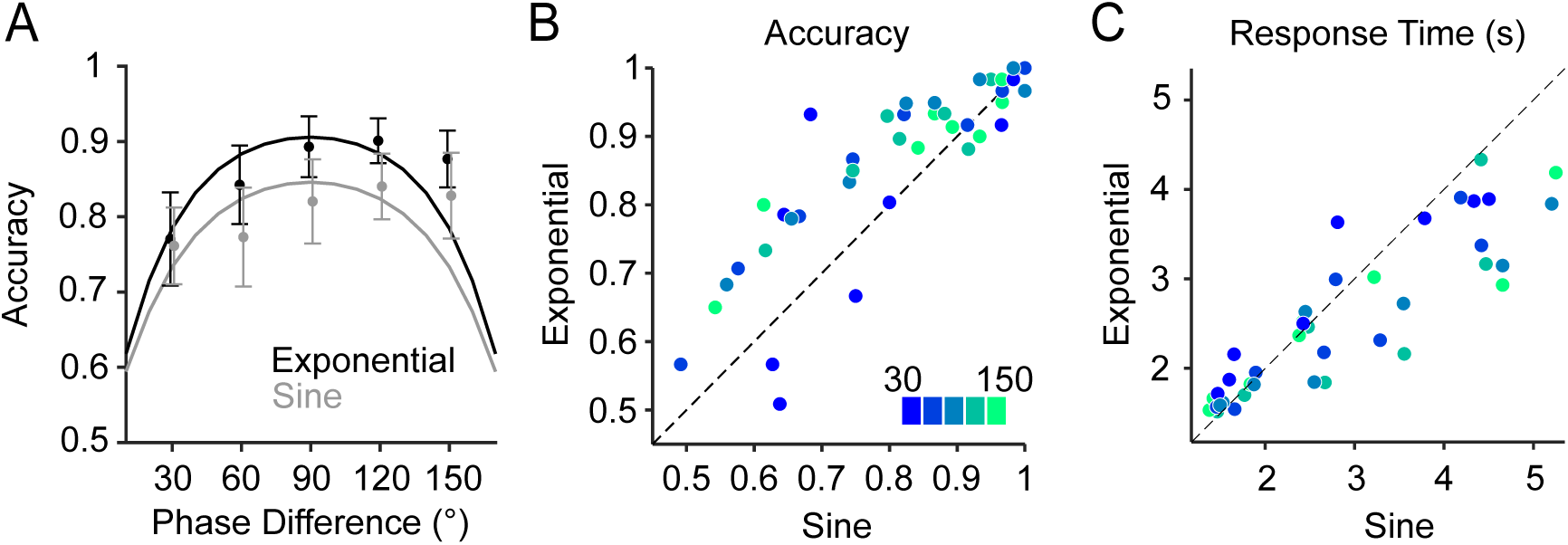
Effect of modulation waveform on tactile motion perception (Experiment 2). (A) Mean motion direction discrimination accuracy (proportion correct) as a function of phase difference for exponential and sinusoidal (labelled “Sine”) envelopes. Markers indicate mean values and error bars represent SEM across participants (*n* = 8). Curves show population-level predictions from the GLMM, obtained by averaging subject-level conditional predictions across participants. (B) Accuracy in the exponential-waveform versus sine-waveform condition. Each data point corresponds to a participant-phase combination and is colour-coded by phase offset (30°, 60°, 90°, 120°, and 150°). The diagonal dashed line indicates equality between conditions. (C) Same as (B), but for response times.

Trial-by-trial accuracy was further analysed using a binomial generalised linear mixed-effects model with a probit link function. Fixed effects included the sinusoidal phase offset sin(*φ*), and its interaction with the modulation waveform (sine versus exponential), with random slopes capturing subject-level sensitivity to phase offset. The phase term was significantly positive (*β* = 1.75 *±* 0.35, 95% CI: [1.08, 2.44], *p <* 10^−6^), indicating a robust relationship between discrimination accuracy and phase difference. Crucially, the interaction term was significantly negative (*β* = *−*0.36 *±* 0.07, 95% CI: [*−*0.48*, −*0.23], *p <* 10^−7^), indicating a higher phase sensitivity for the exponential than for the sine envelope, and the effective phase coefficient was reduced for the sine envelopes (1.40 versus 1.76 in probit units). Model diagnostics indicated an adequate fit to the data (see Figure 2A) with negligible evidence of overdispersion beyond the binomial assumption (dispersion parameter of 1.2, *n* = 4,715 trials). Because modulation envelope waveform entered the model only through its interaction with phase difference, this advantage is neither additive nor present under no motion evidence conditions (e.g. at 0 or 180° phase offsets, see (Adibi, 2025)), but emerges specifically through enhanced contribution of temporal phase information. Accordingly, these results indicate that exponential envelopes selectively enhance the effectiveness of phase-based directional cues rather than producing a global shift in performance.

Response times, as a complementary behavioural measure of task performance, showed a phase-dependent modulation by waveform. At intermediate phase offsets (60–120°), responses were significantly faster for exponential than for sinusoidal envelopes, with reductions of 0.31 *±* 0.17 s at 60°, 0.51 *±* 0.24 s at 90°, and 0.43 *±* 0.22 s at 120°, and with 95% nonparametric bootstrap CIs of differences excluding zero. No systematic response-time difference was observed at 30° (*−*0.09 *±* 0.17 s), whereas the 150° condition showed a numerically similar but more variable reduction for exponential envelopes (0.31 *±* 0.25 s). This pattern suggests that envelope waveform modulates the relationship between phase offset and response time.

To further assess these effects at the trial level while accounting for between-subject variability, response times were analysed using a generalised linear mixed-effects model with an Inverse Gaussian distribution and a log link. Fixed effects included the sine of the phase difference, sin(*φ*), and its interaction with modulation waveform (sine versus exponential), with random intercepts for subjects. Response times decreased significantly with increasing phase difference (*β* = *−*0.138 *±* 0.037, 95% CI: [-0.210, -0.066], *p <* 10^−3^). Importantly, the interaction term was also significant (*β* = 0.063 *±* 0.019, 95% CI: [0.026, 0.100], *p <* 10^−3^), indicating that the envelope waveform modulated the relationship between phase difference and response time, with a weaker phase dependence for sine envelopes than for exponential envelopes.

### Experiment 3: Tactile Motion Across Adjacent and Non-adjacent Digits

While Experiment 1 and 2 demonstrate that temporal coordination of tactile signals supports unified motion percepts across fingers, converging evidence indicates that the spatial arrangement of stimulated sites influences tactile integration (Tamè et al., 2011; Pei et al., 2008; Kuroki and Nishida, 2018). In particular, motion inference may be less effective when phase cues span non-adjacent digits, where intervening digits are not directly stimulated and signals must be integrated across discontinuous representations. In the context of the present study, where motion is inferred from temporal phase offsets between two spatially static inputs, this raises the possibility that motion signals delivered across adjacent fingertips are integrated more efficiently than those spanning a non-adjacent digit. Experiment 3 tests this by quantifying phase-offset thresholds for motion direction discrimination across adjacent and non-adjacent fingertip configurations.

Participants performed motion direction discrimination using three fingertip configurations: index–middle (IM), middle–ring (MR), and index–ring (IR), corresponding to stimulation across adjacent (IM and MR) and non-adjacent (IR) digit pairings (Figure 3A). Phase-offset thresholds were estimated using QUEST staircases constrained to a maximum of 60°. Because threshold estimates were right-censored at this upper bound and thresholds exceeding this bound could not be reliably quantified, performance was characterised using the empirical cumulative distribution functions (ECDFs) of threshold estimates and complementary accuracy measures at 60°.

**Figure 3:**
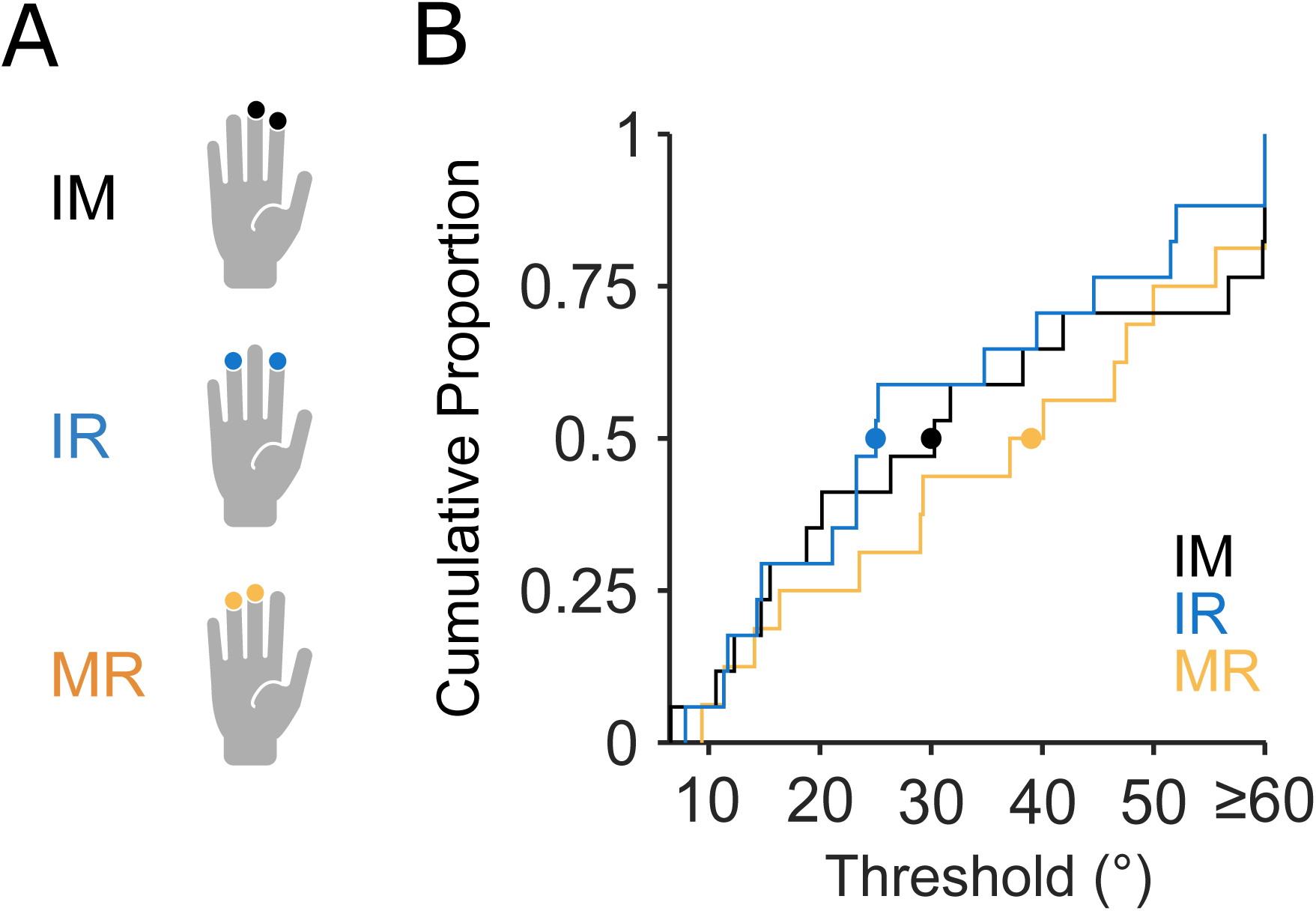
Fingertip stimulation configurations and threshold distributions (Experiment 3). (A) Schematic of the stimulated fingertip pairs. Black indicates index–middle (IM), blue indicates index–ring (IR), and orange indicates middle–ring (MR) stimulation. (B) Empirical cumulative distribution functions (ECDFs) of phase-offset discrimination thresholds for each configuration (IM/IR: *n* = 17; MR: *n* = 16). Colour conventions as in (A). Curves show the cumulative proportion of participants with thresholds at or below a given phase offset; rightward shifts indicate higher thresholds (poorer sensitivity). Filled circles mark the median threshold (ECDF = 0.5) for each condition: MR = 39° (orange), IM = 30° (black), and IR = 25° (blue). The rightmost point (*≥* 60°) denotes thresholds truncated at the maximum tested phase offset.

Comparison of ECDFs revealed a tendency toward higher phase-offset thresholds for the MR configuration relative to the IM and IR (Figure 3B). In contrast, the IM and IR threshold distributions largely overlapped, indicating comparable sensitivity for these two configurations despite their difference in adjacency. These observations were evaluated using within-subject permutation tests on the area between ECDFs. Pairwise comparisons did not reveal statistically significant differences between configurations (MR–IR: *p* = 0.158; MR–IM: *p* = 0.306; IM–IR: *p* = 0.311), indicating that the observed differences in threshold distributions did not reach statistical significance.

Post hoc power analysis indicated that substantially larger sample sizes would be required to detect effects of the observed magnitude (approximately 35 participants for MR–IR, 129 for MR–IM, and 228 for IM–IR pairwise comparisons to achieve 80% power).

To assess whether consistent within-subject trends were present despite non-significant group-level effects, we examined the direction of paired threshold differences across participants. For MR versus IR (*n* = 16), thresholds were higher for MR in 9/16 participants (56.3%), lower in 6/16 (37.5%), and undetermined in 1/16 (6.3%) where both estimated thresholds were above 60°. For MR versus IM (*n* = 16), thresholds were higher for MR in 8/16 (50.0%) and lower in 8/16 (50.0%). For IM versus IR (*n* = 17), thresholds were higher for IM in 8/17 (47.1%), lower in 8/17 (47.1%), and undetermined in 1/17 (5.9%) where both estimated thresholds were censored at 60°.

Median threshold estimates reflected a similar pattern, with comparable medians for IM (30°) and IR (25°), and a higher median for MR (39°), as shown in Figure 3B. However, because threshold estimates were right-censored at 60°, the MR median should be interpreted as a lower bound rather than a precise estimate.

Because threshold estimates were constrained by the 60° upper limit, discrimination accuracy was analysed at this maximum tested phase offset, as a complementary measure. Accuracy was highest for IR (89.8%*±*3.1%), intermediate for MR (87.0%*±*3.9%), and lowest for IM (78.1%*±*5.1%).

A binomial generalised linear mixed-effects model (probit link), with fingertip configuration as a fixed effect and subject as a random intercept to account for subject variability, revealed no significant difference between discrimination accuracy at 60° for IR versus MR (*β* = 0.13 *±* 0.18, 95% CI: [*−*0.21, 0.48], *p* = 0.45). In contrast, discrimination accuracy at 60° was significantly lower for IM than for MR (*β* = *−*0.32 *±* 0.14, 95% CI: [*−*0.59*, −*0.05], *p* = 0.021). Model diagnostics indicated an adequate fit, with no evidence of overdispersion relative to the binomial assumptions (dispersion parameter: 0.97; *n* = 682 trials).

Consistent with the model fit, marginal (population-level) predictions obtained by Monte Carlo integration over the estimated distribution of subject-level random effects reproduced the observed ordering across configurations. At 60°, predicted accuracy was highest for IR (85%, 95% CI: [75.7, 91.5]), followed by MR (81.7%, 95% CI: [74.4, 87.7]), and lowest for IM (72%, 95% CI: [62.4, 80.3]), closely matching the pattern observed in the empirical data.

Together, these results indicate that the fingertip configuration had only a limited and inconsistent effect on phase-based motion sensitivity under the tested conditions.

### Experiment 4: Bimanual versus Unimanual Integration of Temporal Phase Cues

Perceptual sensitivity to temporal order and spatiotemporal structure in touch differs between unimanual and bimanual stimulation. Classic studies of tactile temporal-order judgement have shown that cross-hand temporal discrimination can be more precise than within-hand discrimination, indicating enhanced temporal resolution for stimulation distributed across the two hands (Hirsh and Sherrick, 1961; Sherrick and Rogers, 1966; Craig, 1985). Consistent with this, tactile temporal judgements and motion integration can be improved when stimuli are distributed across hands rather than within a single hand (Shore et al., 2005; Gallace and Spence, 2005; Arslanova et al., 2022). This body of work predicts that temporal phase information should be utilised more effectively under bimanual than unimanual stimulation, leading to improved motion discrimination performance. Experiment 4 tests this prediction by comparing motion direction discrimination under bimanual and unimanual stimulation of phase-offset vibrotactile signals.

Across participants and phase offsets, bimanual stimulation resulted in higher accuracy than unimanual stimulation, exhibiting an overall enhancement of 6.0% (Figure 4A and B). The bimanual enhancement was small at the smallest phase offset (30°; 3.7% *±* 3.2%). but increased at larger phase offsets, reaching +8.1% *±* 1.5% at 60° and +6.2% *±* 2.2% at 90°, where the effects were statistically significant (both 95% nonparametric bootstrap CIs excluded zero). These results revealed that distributing temporally offset tactile signals across the two hands selectively enhanced motion discrimination at phase offsets where temporal information was most informative.

This pattern was confirmed by a hierarchical trial-level analysis of accuracy using a binomial GLMM with a probit link. Accuracy was modelled as a function of sin(*φ*) and its interaction with hand configuration (unimanual versus bimanual), with subject-specific random slopes for sin(*φ*) accounting for cross-subject variability in phase sensitivity. The sin(*φ*) term was significantly positive (*β* = 0.863 *±* 0.183, 95% CI: [0.50, 1.22], *p <* 10^−5^), indicating that discrimination accuracy increased with sin(*φ*). The interaction between sin(*φ*) and hand configuration was also significant (*β* = 0.312 *±* 0.050, 95% CI: [0.22, 0.41], *p <* 10^−9^), demonstrating higher phase sensitivity in the bimanual condition than in the unimanual condition. As hand configuration entered the model only through its interaction with sin(*φ*), the bimanual effect was phase-dependent: it was absent at zero phase evidence and emerged progressively as sin(*φ*) increased. Model diagnostics indicated an adequate fit (see Figure 4A) with no evidence of overdispersion relative to the binomial assumption (dispersion parameter of 1.01, *n* = 5,636).

**Figure 4:**
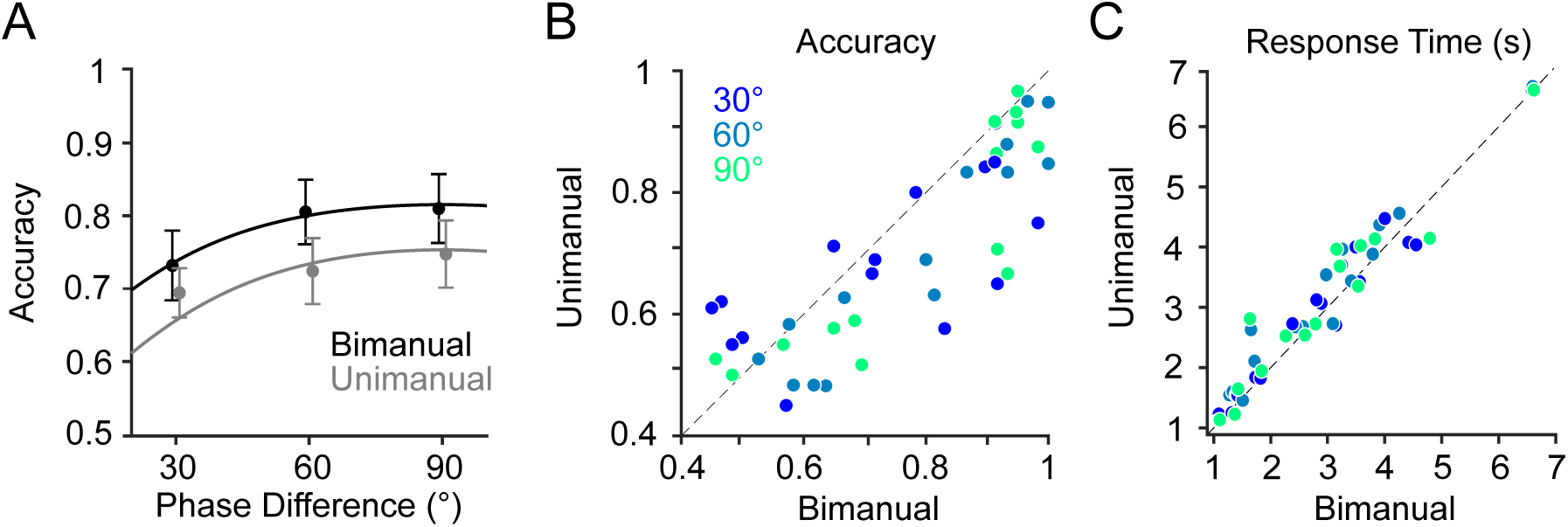
Bimanual versus unimanual tactile motion discrimination (Experiment 4). (A) Mean motion direction discrimination accuracy (proportion correct) as a function of phase difference for bimanual and unimanual stimulation conditions. Markers indicate mean values and error bars represent SEM across participants (n=16). Curves show population-level GLMM predictions obtained by averaging subject-specific conditional predictions across participants. (B) Accuracy in bimanual condition versus unimanual condition. Each data point corresponds to a participant-phase combination and is colour-coded by phase difference (30°, 60°, and 90°). The diagonal dashed line indicates equality between conditions. (C) Same as (B), but for response times.

Across subjects and phase differences, response times for bimanual stimulation were on average 0.17*±*0.08 s (mean *±* SEM across participants) faster than those of unimanual stimulation, in a phase-dependent manner (Figure 4C). At 30°, mean response times were comparable for bimanual and unimanual stimulation (3.03*±*0.36 s versus 3.10*±*0.35 s). At 60°, responses were faster by 0.26*±*0.08 s in the bimanual condition (2.80*±*0.36 s) than in the unimanual condition (3.06*±*0.36 s), with 95% nonparametric bootstrap CI of the difference across participants excluding zero. At 90°, a similar effect was observed (0.18*±*0.11 s).

This pattern was supported by a hierarchical trial-level analysis of response times using a Gamma GLMM with a log link. Response times were modelled as a function of sin(*φ*) and its interaction with hand configuration (unimanual versus bimanual), with subject-specific random intercepts accounting for cross-participant variability. The sin(*φ*) term was significantly negative (*β* = *−*0.076 *±* 0.032, 95% CI: [*−*0.139*, −*0.013], *p* = 0.017), indicating that response times decreased with increasing sin(*φ*). The interaction between sin(*φ*) and hand configuration was also significant (*β* = *−*0.089 *±* 0.016, 95% CI: [*−*0.120*, −*0.057], *p <* 10^−7^), demonstrating stronger phase-dependent decreases in response time under bimanual than under unimanual stimulation. Because hand configuration entered the model only through its interaction with sin(*φ*), the bimanual effect was phase-dependent, progressively emerging as sin(*φ*) increased. Model diagnostics indicated an adequate fit, with a dispersion parameter of 0.243, indicating that response time variability scaled appropriately with the mean, consistent with the assumptions of the Gamma model (*n* = 5,636 trials).

## Discussion

The present study examined how vibrotactile motion direction perception depends on systematic manipulations of stimulus parameters across several spatial configurations. Across experiments, directional motion percepts were reliably evoked by phase offsets between amplitude-modulated vibrations delivered to static skin locations, confirming that tactile motion can emerge from temporal relationships alone, without physical translation across the skin. This finding is consistent with classic demonstrations of apparent tactile motion and saltatory illusions, in which spatial percepts are constructed from temporally structured stimulation rather than continuous movement (Hirsh and Sherrick, 1961; Sherrick and Rogers, 1966; Geldard and Sherrick, 1972).

Modulation amplitude mainly influenced response time rather than discrimination accuracy. Increasing amplitude levels did not systematically affect discrimination accuracy, while response times were shorter at higher amplitudes. This dissociation indicates that stronger stimulation reduces response times without proportionally improving directional discrimination. Consistently, studies in humans and rodents have revealed that reaction times in tactile detection and discrimination tasks decrease with stimulus intensity (Gescheider et al., 1969; Adibi and Arabzadeh, 2011; Ollerenshaw et al., 2012).

This result aligns with predictions from both correlation-based and landmark-based models (Figure 5A-C), which do not rely on absolute stimulus amplitude once reliable temporal information is available. Correlation-based models are invariant to amplitude changes, as the correlation between two signals remains unchanged under scaling.

**Figure 5:**
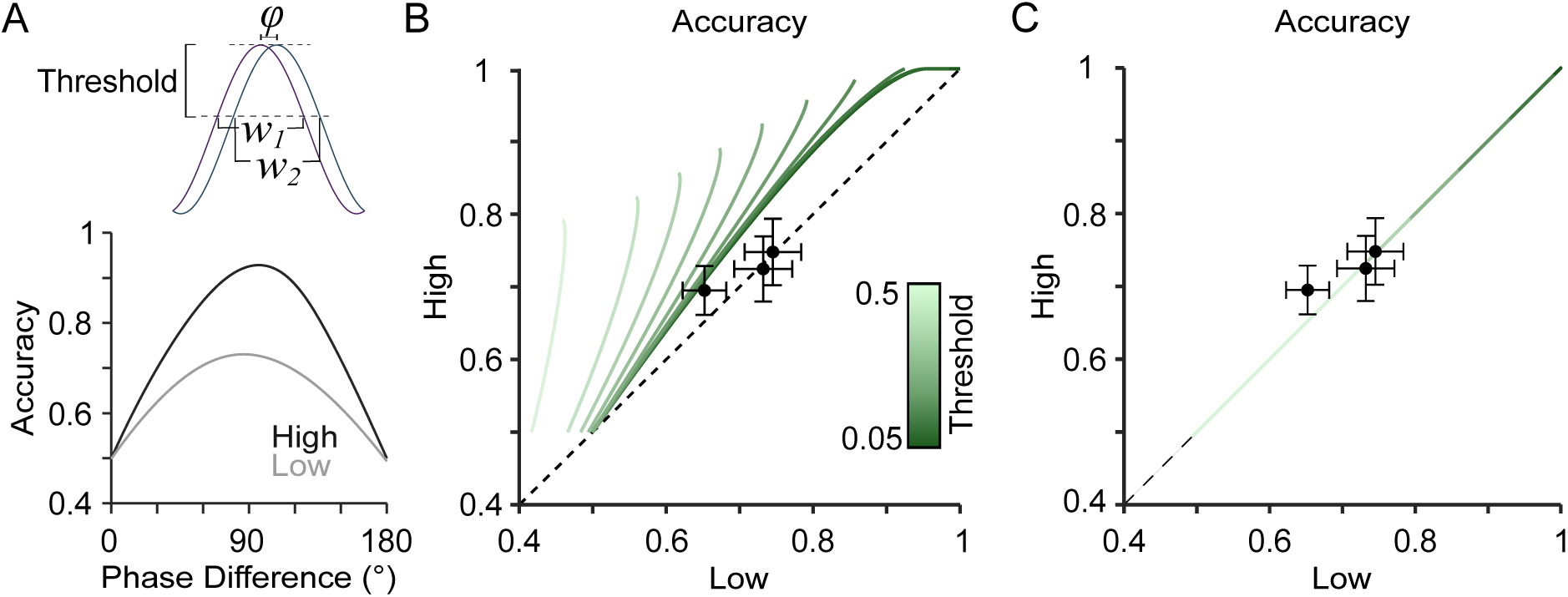
Landmark model predictions for the effect of modulation amplitude on tactile motion perception (Experiment 1). (A) Landmark model illustration and corresponding prediction. Top: One cycle of two amplitude-modulated vibration envelopes with a phase offset of 30°. The horizontal line indicates the amplitude discrimination threshold (0.3 relative to the peak). The intersection points of this threshold with each envelope (*w*_1_ and *w*_2_) define the temporal window around each landmark (peak), representing uncertainty in landmark timing. The overlap between these windows (77.4% for 30°) determines the probability of correct motion discrimination (see (Adibi, 2025)). Bottom: Model-predicted discrimination accuracy as a function of phase difference for the same threshold (0.3), shown for low- and high-amplitude conditions in grey and black, respectively. (B) Model-predicted discrimination accuracy for the fixed-threshold model, comparing high- and low-amplitude conditions. Each curve corresponds to a different threshold value (colour-coded) and shows predicted performance across phase differences (0–90°). Markers and error bars indicate empirical mean *±* SEM across participants. The diagonal dashed line indicates equal performance in the two conditions. (C) same as in (B), but for the Weber-scaling model, in which the detection threshold scales with amplitude.

In the landmark framework (Adibi, 2025), amplitude effects depend on the assumed feature-detection criterion: under a fixed absolute threshold, higher amplitude narrows the above-threshold timing window and can modestly improve predicted accuracy (Figure 5B), whereas under Weber-like scaling, amplitude discrimination thresholds increase proportion-ally with intensity (Adibi and Arabzadeh, 2011; Francisco et al., 2008; Gescheider, 2013), preserving the timing window and resulting in approximately equal predicted accuracy across amplitudes (Figure 5C). Together, these results indicate that both model classes predict largely amplitude-invariant motion discrimination once timing cues are reliably available, with residual amplitude dependence arising primarily under a fixed absolute detection criterion.

The manipulation of modulation envelope waveform provides a more selective test of the two competing computational accounts (correlation-based versus landmark). Although both sinusoidal and exponential envelopes support robust motion perception (Adibi, 2025), exponential envelopes produced a modest but reliable improvement in discrimination accuracy. Exponential envelopes contain steeper temporal gradients and more temporally localised features (e.g., peaks) than sinusoidal modulation (Figure 6A), which may improve the temporal precision with which salient envelope features are detected.

**Figure 6:**
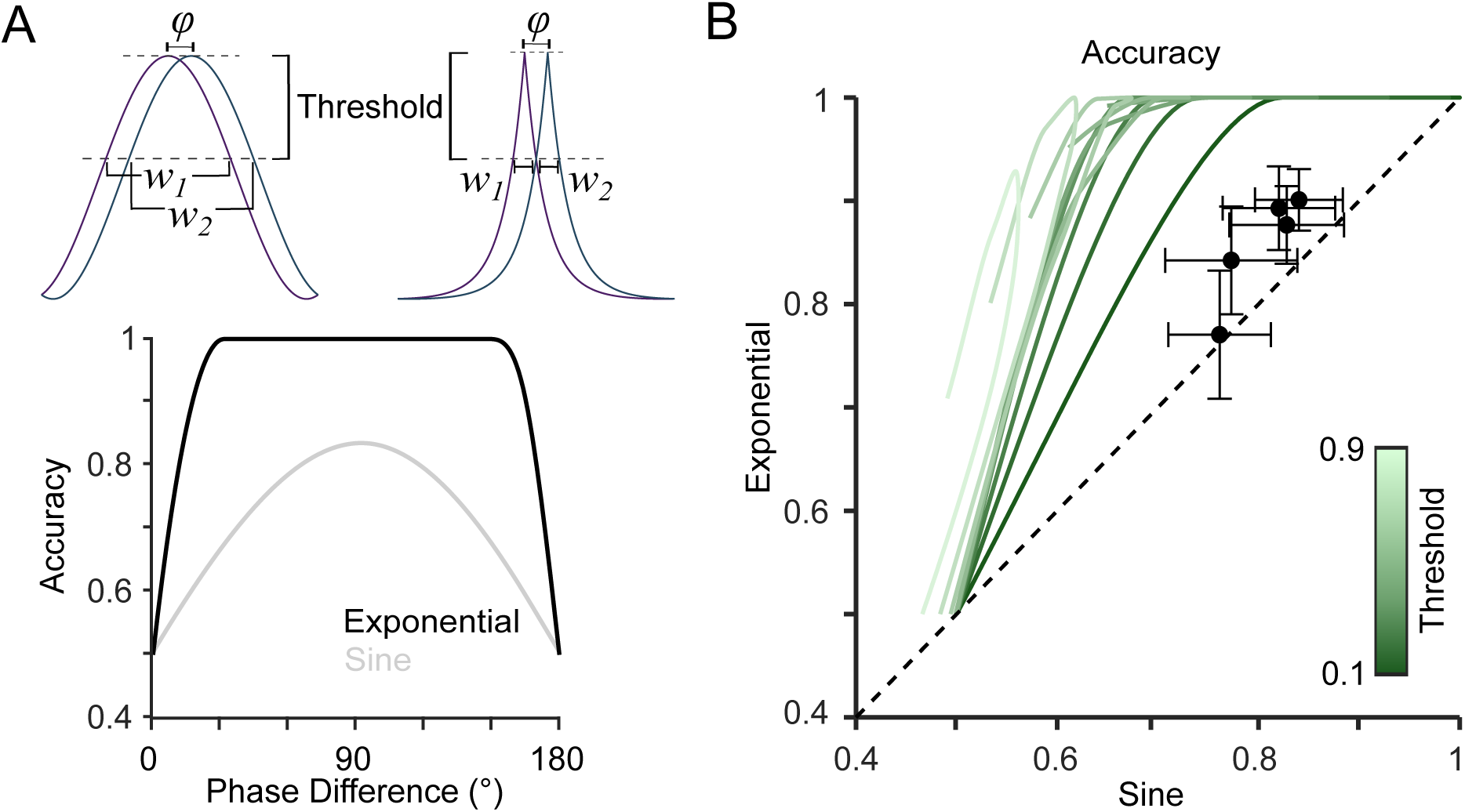
Landmark model predictions for the effect of modulation waveform on tactile motion perception (Experiment 2). (A) Landmark model illustration and corresponding prediction. Top: One cycle of two amplitude-modulated vibration envelopes (sinusoidal and exponential) with a phase offset of 30°. The horizontal line indicates the amplitude discrimination threshold (0.44). The intersection points of this threshold with each envelope define the temporal windows around each landmark (peak), representing uncertainty in landmark timing (*w*_1_ and *w*_2_). The overlap between these windows determines directional accuracy, and is larger for the sinusoidal envelope than for the exponential envelope. Bottom: Model-predicted discrimination accuracy as a function of phase difference for a fixed threshold, showing higher predicted accuracy for the exponential than for the sinusoidal waveform. (B) Predicted discrimination accuracy for varying absolute thresholds, comparing exponential versus sinusoidal waveforms. Curves correspond to different threshold values (colour-coded), and markers with error bars indicate empirical mean *±* SEM across participants. The diagonal dashed line indicates equality between conditions.

The landmark framework predicts this effect because exponential envelopes produce narrower above-threshold timing windows, reducing uncertainty in landmark timing and thereby improving temporal-order discrimination. Consistent with this account, model predictions capture the advantage of exponential over sinusoidal envelopes, with the size of the predicted effect varying with the assumed threshold level (Figure 6B). Thus, the waveform manipulation argues against strict waveform invariance and supports the contribution of temporally localised features to tactile motion computation. More generally, waveform shape can influence tactile sensitivity and response dynamics (Young et al., 2015). In particular, sharper waveforms often produce lower detection thresholds and faster responses than smoother (sinusoidal) stimulation (Chancey et al., 2014; Vardar et al., 2017), supporting the view that envelope waveform modulates – rather than determines – phase-based motion perception under the tested conditions.

The landmark model therefore captures the direction, but not the exact magnitude of the waveform effect. In particular, this single-parameter model predicts a larger separation between exponential and sinusoidal envelopes than was observed behaviourally. One likely contributor is that the model defines landmarks directly from the stimulus envelope, whereas the tactile signal reaching mechanoreceptors is reshaped by actuator dynamics, fingertip contact, and viscoelastic skin mechanics before neural encoding (Serhat and Kuchenbecker, 2024; Tummala et al., 2026; Dandu et al., 2021). Such biomechanical filtering may reduce the sharp temporal transients that give exponential envelopes their predicted advantage.

Evidence also suggests that tactile vibration coding is unlikely to depend on envelope amplitude alone. Behavioural studies indicate that vibrotactile perception depends on combined amplitude-frequency information, including velocity (amplitude *×* frequency) (Adibi et al., 2012) and similar power-law amplitude-frequency relationships underlying vibrotactile pitch perception (Prsa et al., 2021). These findings are further supported by recent evidence that Pacinian mechanotransduction is velocity-sensitive (Chikamoto et al., 2026), suggesting that more realistic implementations of the landmark framework may require landmark definitions based on instantaneous velocity or energy (Kuroki et al., 2016; Kuroki and Nishida, 2018) rather than amplitude alone.

Longer response times for sinusoidal envelopes may also partially compensate for weaker temporal landmarks by allowing additional time for evidence accumulation, thereby improving discrimination accuracy. This is consistent with speed-accuracy trade-offs (Wickelgren, 1977; Heitz, 2014; McDonald et al., 2014) and evidence-accumulation accounts in which accuracy and response time jointly reflect latent decision processes rather than sensory strength alone (Ratcliff, 1978; Ratcliff and McKoon, 2008; Mulder and van Maanen, 2013).

We further quantified motion direction discrimination sensitivity across adjacent and non-adjacent finger pairings. Sensitivity for index-ring (IR) and index-middle (IM) configurations was comparable, while the middle-ring (MR) configuration showed a tendency toward higher thresholds (lower sensitivity). However, these differences were not statistically significant at the group level, indicating that any effect of fingertip configuration on phase-offset sensitivity was limited under the tested conditions. A single comparison in the GLMM showed lower accuracy for IM relative to MR at 60°, but this effect was not reflected in the threshold distributions and should therefore be interpreted with caution.

A distance-based account would predict lower sensitivity for adjacent digit pairings due to increased within-hand interference arising from overlapping somatotopic representations in early somatosensory processing areas (Tamè et al., 2011, 2014). This account is also consistent with the improved performance observed under bimanual stimulation, where inputs are distributed across hands and thus less affected by within-hand interference(Craig, 1985) and more effectively integrated across hands (Arslanova et al., 2022). However, an alternative possibility is that spatial proximity facilitates the integration of temporally structured tactile signals by supporting a more continuous percept of motion across the skin, consistent with classic findings in tactile apparent motion (Sherrick and Rogers, 1966; Kirman, 1974) and with sensitivity to inter-finger phase relationships in vibrotactile stimulation (Kuroki and Nishida, 2018). Under this view, closer stimulation sites may enhance the perceptual coherence of motion signals, analogous to spatial constraints observed in visual apparent motion (Kolers, 1972; Braddick, 1974). The absence of a consistent adjacency-related effect in the present data may therefore reflect the interaction of these competing mechanisms, with potential benefits of spatial continuity offsetting interference arising from somatotopic proximity.

An alternative account is that performance depends not only on adjacency-based factors, but also on digit identity; that is, differences in how signals from specific digits are represented and integrated. In the present data, IM and IR showed highly overlapping threshold distributions despite differing in adjacency, whereas MR showed a tendency toward higher thresholds relative to the index-involving pairings (IM and IR). This pattern is more consistent with the contribution of digit identity than with spatial separation alone. This interpretation is supported by evidence that tactile acuity varies across fingers and is linked to differences in cortical magnification in primary somatosensory cortex (Duncan and Boynton, 2007), as well as evidence that individual digits are represented in distinct, spatially organised cortical maps across somatosensory cortex (areas 3b, 1, 2) (Martuzzi et al., 2014). These findings suggest that variability in performance may reflect differences in the size and organisation of digit-specific cortical representations, and consequently in how signals from different digits are integrated.

The digit identity account may also contribute to the bimanual advantage over the unimanual condition. In the bimanual condition, stimulation involved two homologous index fingers, whereas the unimanual condition involved index and middle fingers. Engaging two index-finger representations may therefore provide a modest benefit, consistent with digit-specific differences in tactile acuity and cortical organisation (Duncan and Boynton, 2007; Martuzzi et al., 2014). At the same time, bimanual stimulation reduces within-hand competition and engages distributed processing across hemispheres, factors that have been linked to improved tactile temporal and multi-digit performance (Craig, 1985; Iida et al., 2016; Arslanova et al., 2022). Together, these factors suggest that the bimanual benefit may reflect a combination of cross-hand integration and digit identity differences in tactile processing.

While behavioural data alone cannot identify the underlying neural mechanisms, the observed pattern is consistent with bilateral integration at multiple stages of the somatosensory-parietal network. Higher-order somatosensory regions, including SII, exhibit comparatively bilateral receptive fields and interhemispheric coupling during tactile object processing (Yu et al., 2018), supporting their role in combining tactile information across the body midline. In parallel, neurophysiological studies indicate digit-specific cross-hand influences on SI responses (Tamè et al., 2015), suggesting that bilateral interactions can modulate sensory encoding even at early processing stages. Together, these findings are consistent with the possibility that distributing temporally structured cues across the two hands does not impose a processing cost and may facilitate the integration of task-relevant information.

In our paradigm, tactile motion judgements may arise from the accumulation of sensory evidence over time, with evidence derived either from correlation-based integration of temporally offset inputs (Pei et al., 2011) or from landmark-based mechanisms that compare the relative timing of salient events (Craig and Busey, 2003). In a sequential-sampling framework, pooling information across cycles – and potentially across hands – could increase the effective quality of the evidence entering a decision variable, for example by increasing the rate of accumulation and/or reducing decision noise. This account is consistent with human EEG studies showing a centroparietal decision signal whose buildup tracks evidence accumulation toward a bound (O’Connell et al., 2012; Kelly and O’Connell, 2013; Twomey et al., 2015), and with single-unit and computational studies implicating posterior parietal cortex in perceptual decision-making (Shadlen and Newsome, 2001; Gold and Shadlen, 2007). Joint analysis of response time distributions and choice accuracy within a drift–diffusion framework could provide further insight into the temporal structure of the sensory evidence underlying phase-based tactile motion perception (Ratcliff, 1978; Ratcliff and McKoon, 2008; Shinn et al., 2020). Although the number of trials per condition in the present study was not sufficient to reliably characterise response time distributions, applying such models in future work could reveal how temporal phase differences shape the rate and reliability of evidence accumulation, and whether bimanual stimulation enhances performance by increasing drift rate or reducing decision noise.

Future electrophysiology studies could therefore test whether bimanual stimulation is associated with a steeper buildup or earlier threshold crossing of decision-related signals, consistent with more efficient integration of bilateral tactile evidence.

